# Quantifying the monomer-dimer equilibrium of tubulin with mass photometry

**DOI:** 10.1101/2020.07.16.199083

**Authors:** Adam Fineberg, Thomas Surrey, Philipp Kukura

## Abstract

The *αβ*-tubulin heterodimer is the fundamental building block of microtubules, making it central to several cellular processes. Despite the apparent simplicity of heterodimerisation, the associated energetics and kinetics remain disputed, largely due to experimental challenges associated with quantifying affinities in the <μM range. We use mass photometry to observe tubulin monomers and heterodimers in solution simultaneously, thereby quantifying the *αβ*-tubulin dissociation constant (8.48±1.22 nM) and its tightening in the presence of GTP (3.69±0.65 nM), at a dissociation rate >10^−2^ s^−1^. Our results demonstrate the capabilities of mass photometry for quantifying protein-protein interactions and clarify the energetics and kinetics of tubulin heterodimerisation.

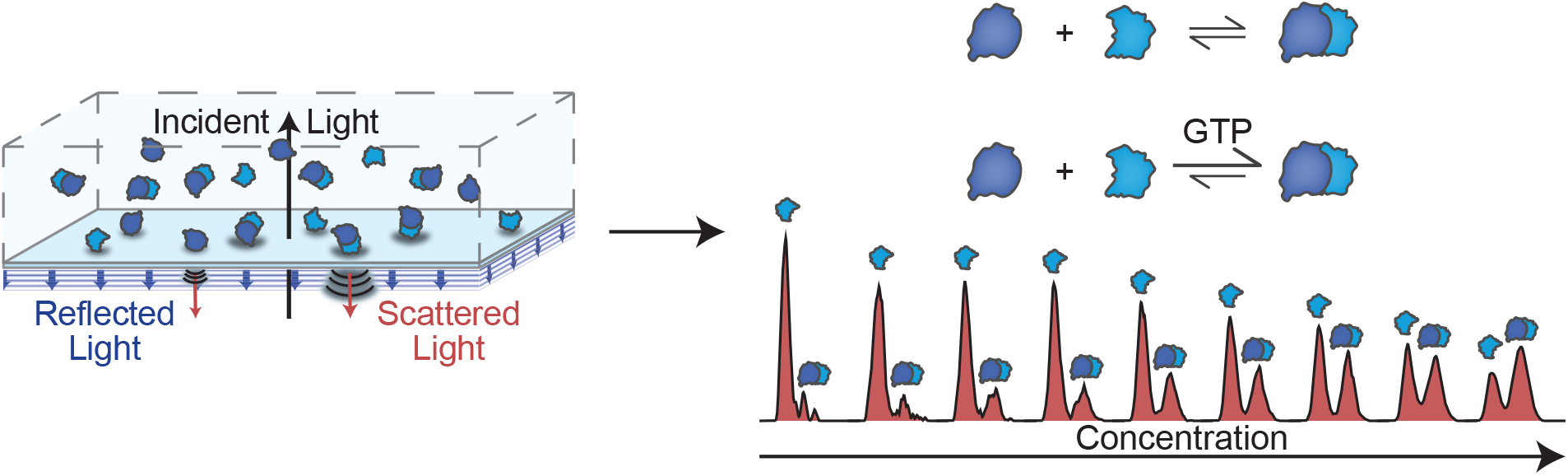

---

Microtubules, involved in processes as broad as mitosis, cell motility and intracellular transport, are constructed of heterodimers of *α*- and *β*-tubulin — highly conserved members of the tubulin/FtsZ family of proteins — with each subunit able to bind a GTP molecule.[1] Heterodimer formation, the first critical step towards microtubule assembly, has been reported to require a number of cofactors.[2, 3, 4] Studies on the thermodynamic and kinetic stability of *αβ*-tubulin heterodimers, which ultimately impact the assembly and stability of microtubules, have produced a broad range of binding affinities and kinetics, ranging over 5 orders of magnitude from 10^−11^ to 10^−6^ M, with dissociation rates from 10^−5^ to 10^−2^ s^−1^.[5, 6, 7, 8]

These contradictory and inconsistent results can be ultimately attributed to a combination of factors including tissue source and sample preparation,[9] as well as the lack of experimental approaches capable of quantifying binding affinities in the sub-μM range in near-native conditions. Truly label-free approaches generally require μM concentrations or higher, with higher dilutions only accessible through labelling, surface-based methods, or a combination of both. We have recently introduced mass photometry (MP), single molecule detection and mass measurement in solution based on light scattering.[10] MP uses the interference between light scattered by a biomolecule as it non-specifically binds to a glass surface and the reflection of the illumination light from the glass-water interface to produce label-free images of single biomolecules (Fig. 1a). The resulting optical contrast scales linearly with molecular mass, enabling the identification and counting of molecules and their complexes in solution. Detection is label-free, with the surface acting only as a detector and all interactions taking place in free solution between unmodified molecules over a concentration range from 0.1 to 100 nM, making MP in principle ideally suited to study tight protein-protein interactions in a quantitative fashion.[11, 12, 13]

**Figure 1:**
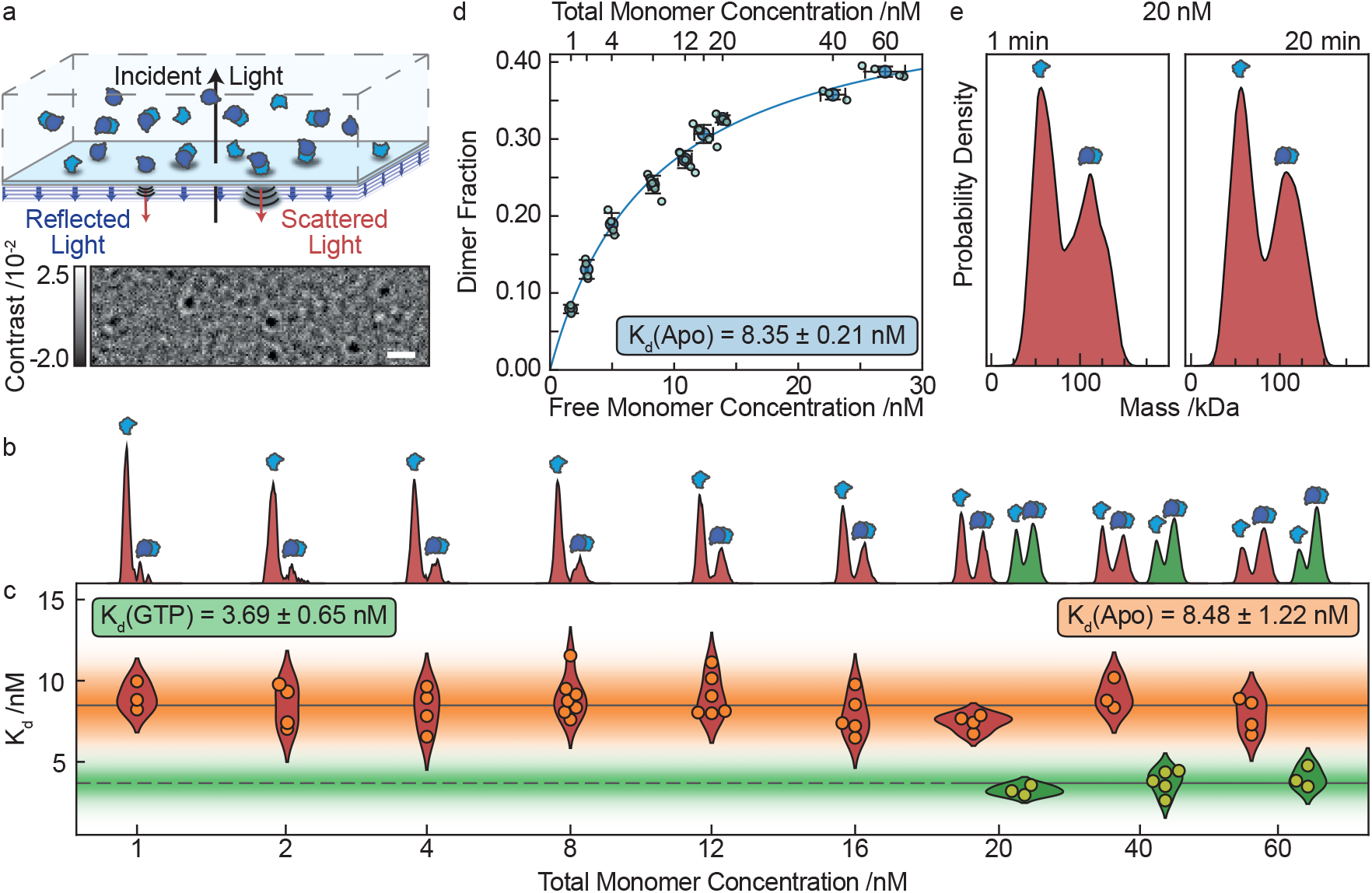
Quantification of tubulin heterodimersiation with mass photometry. **a,** Schematic illustrating the operation mass photometry. Imaging the interference between scattered and reflected light as a protein non-specifically binds at a glass-water interface results in label-free single molecule images with a contrast proportional to their molecular mass. Scale bar: 1 μm. **b,** Mass kernel density estimates with 5 kDa bandwidth for tubulin at total monomer concentrations ranging from 1 – 60 nM in the apo (red) and 20 – 60 nM in the GTP-bound (green) state. **c,** Resulting binding affinities extracted from each distribution in **b**. The grey lines indicate the global mean *K_d_* and the shaded areas the standard deviations for each data set. **d,** Proportion of dimer present in the apo state at equilibrium as a function of free monomer concentration at equilibrium. Light blue markers indicate individual experiments, dark blue markers and error bars depict the mean and standard deviation respectively for each total monomer concentration. A logistic fit through the mean values yields *K_d_* = 8.35 ± 0.21 nM (error on fit). **e,** Averaged mass kernel density estimates (N = 4) with 5 kDa bandwidth for tubulin in the apo state at a total monomer concentration of 20 nM from separate experiments 30 s after (left) and 20 min after dilution (right).

Applying MP to tubulin purified from porcine brain diluted at 60 nM concentration in BRB80 buffer exhibited a roughly 2:1 dimer:monomer ratio in the apo state, indicative of a low nM *K_d_*. Accordingly, repeating these measurements at total tubulin concentrations ranging from 1 to 60 nM revealed a transition from predominantly monomeric towards predominantly dimeric distributions (Fig. 1b). We can convert these distributions into binding affinities in multiple ways. Given that we are directly counting monomers and dimers, we can compute a *K_d_* from any one of these distributions given knowledge of the total protein concentration. The results are consistent across all concentrations measured yielding *K_d_* = 8.48 ± 1.22 nM (Fig. 1c), in close agreement with a more traditional titration-based analysis, which requires multiple measurements to yield the affinity of interest (Fig. 1d). We did not find significant differences between these measurements, performed after 20 min of equilibration after dilution, and those performed immediately after dilution (Fig. 1e). During our measurements, which take 60 – 120 seconds, we could also not find any evidence of dissociation, suggesting that the associated off-rates must be faster than 10^−2^ s^−1^. Very slow dissociation rates would reveal identical monomer:dimer distributions for all dilutions, because equilibrium would not be reached during the 20 min between dilution and measurement, and thus reflect the pre-dilution distribution. Incubation with 1 mM GTP, by contrast, resulted in a clear shift towards dimer for a given total monomer concentration (Fig. 1c) and an associated *K_d_* = 3.69 ± 0.65 nM. Due to the decrease in *K_d_* upon GTP binding, achieving an accurate measurement at low concentrations became impossible due to low statistics in the monomer population. However, due to the close agreement of measured apo state *K_d_* values between the binding curve (Fig. 1d) and the single shot (Fig 1c) measurements, we are confident in the accuracy of single shot measurements with MP.

These results quantify the binding affinity for the tubulin heterodimer both in the apo and GTP bound state and provide an upper limit to the dissociation rate, which is orders of magnitude faster than reports based on surface plasmon resonance.[7] Although it has been shown that the *K_d_* of the tubulin dimer depends significantly on the protein’s source,[9] due to the fact that tubulin is highly conserved one would expect the effect of GTP binding to be similar across different species. The reduction of the *K_d_* in the GTP bound state demonstrates the stabilising effect nucleotide has upon the dimer, which in light of the fast off rate suggests improved resistance to local fluctuations in tubulin concentration. This small change in binding affinity corresponds to a ΔΔ*G* ≈ 2 kJ mol^−1^, highlighting the ability of MP to detect and precisely quantify even very subtle changes in protein-protein interactions, which opens the door towards investigating the effects of post-translational modifications. Moreover, the *K_d_* being on the order of nM demonstrates that, under physiological GTP and tubulin concentrations, tubulin will almost exclusively be found as a heterodimer — an important consideration as free *β*-tubulin is known to be toxic.[14]

As *α*- and *β*-tubulin each bind a molecule of GTP, upon dimerisation a GTP molecule becomes buried in the intra-dimer site. Therefore, the stabilising effect that GTP has upon the dimer can be seen to parallel the effect that GTP binding to the inter-dimer site has upon microtubule polymerisation. This leads to the possibility that the intradimer GTP stabilises the dimer whilst the interdimer GTP affects microtubule dynamics, a hypothesis that can be explored in the future through the use of GTP analogues and mutation of the GTP binding sites.

## Acknowledgements

We thank the Surrey lab members, The Francis Crick Institute, for tubulin purification including Nicholas Cade for his assistance in early experiments. Research in the Kukura group is supported by the ERC (CoG 819593), TS was supported by the Francis Crick Institute, which receives its core funding from Cancer Research UK (FC001163), the UK Medical Research Council (FC001163), and the Wellcome Trust (FC001163). T.S. also acknowledges support from the European Research Council (Advanced Grant, project 323042) and from the Spanish Ministry of Economy, Industry and Competitiveness to the CRG-EMBL partnership, the Centro de Excelencia Severo Ochoa and the CERCA Programme of the Generalitat de Catalunya.

## Author Contributions

AF: Formal Analysis, Methodology, Software, Validation, Visualisation, Writing – original draft, review, and editing. TS: Resources, Writing – review and editing. PK: Conceptualisation, Funding acquisition, Methodology, Supervision, Visualisation, Writing – original draft, review, and editing.

