## Supplementary Methods for "Quantifying the monomer-dimer equilibrium of tubulin with mass photometry"

#### Tubulin Purification

Tubulin was purified from porcine brain as described[1]. 1-3 ml aliquots were flash-frozen in liquid nitrogen on the day of purification and stored at −80 °C. An additional polymerisation/depolymerisation cycle was performed before flash freezing and storage in µl sized aliquots in liquid nitrogen. The final storage buffer was BRB80. The small tubulin aliquots were thawed just before the experiment.

#### Mass Photometry

An in-house experimental mass photometry setup was used as described[2]. Briefly, the setup used a 445 nm diode laser (Lasertack) and a Point Grey (GS3-U3-23S6M-C) CMOS camera, as well as the following acquisition parameters: frame rate = 1000 Hz, pixel binning = 3 $\times$ 3, and 5-fold time averaging. Data was acquired for either 60 or 120 s by in-house software implemented with LabView 2015.

#### Sample Delivery

Samples were measured in either a silicone gasket (3 mm diameter, 1 mm thickness) fixed to a microscope coverslip (#1.5, 24 mm $\times$ 52 mm, Menzel Gläser) or a flow chamber. Flow chambers were constructed by making a narrow channel with double sided tape sandwiched between two microscope coverslips (#1.5, 24 mm $\times$ 52 mm, Menzel Gläser and #1.5, 24 mm $\times$ 24 mm, Menzel Gläser). Sample was introduced to the chamber via capillary action and measured at 20 °C.

#### Sample Preparation

A sample of 66 µM tubulin was diluted in Brinkley BR buffer 1980 (BRB80) (80 mM K-PIPES, 1 mM $MgCl_2$, 1 mM EGTA, titrated to pH 6.8 with KOH) to concentrations in the range of 1 to 60 nM total monomer concentration, $[M]_{TOT}$. Unless otherwise stated, all dilutions were left on ice to equilibrate for 20 min. 15 µl of the equilibrated samples were added either to a gasket or flow chamber and allowed to bind non-specifically to the glass surface. 15 s after sample addition, the acquisition software was triggered, and the sample was measured. For the GTP bound assay, the sample was equilibrated for 15 min after dilution and GTP was added with a final concentration of 1 mM. The sample was added to the chamber or gasket after a further 5 min incubation.

### Analysis

#### Image Analysis

Movies were analysed using commercial software from Refeyn Ltd. (Discover MP), after a further 10-fold time averaging with thresholds of 1.00 and 0.25, the remaining parameters were left at their defaults. The returned contrast signals were converted into mass through a calibration using a known mass standard. Using this, the number of monomer and dimer events detected was calculated. Data was presented as kernel density estimates with a 5 kDa bandwidth.

#### Data Analysis

Data was processed using in-house python scripts. The number of monomers, $N_M$, and dimers, $N_D$, detected were converted into effective concentrations of each species as follows:
For the equilibrium between monomer (M) and dimer (D), $M + M \rightleftharpoons D$, the dissociation constant $K_d$ is defined:

$$K_d = \frac{[M]^2}{[D]} \tag{1}$$

The total monomer concentration, $[M]_{TOT}$, in the system is:

$$[M]_{TOT} = 2[D] + [M] \tag{2}$$

Assuming that the number of species detected is proportional the concentration of species, $[M]_{TOT}$ can be expressed as:

$$[M]_{TOT} = c(2N_D + N_M) \tag{3}$$

This assumption requires that the binding events captured in the experiments are representative of the equilibrium populations of monomer and dimer in solution.[2] For the assumption to hold, the rate of diffusion of a species from bulk solution to the glass surface, and the probability of non-specific binding to that surface, where the measurement is made, must be equal for both monomer and dimer. We can determine these parameters by quantifying the rate of decay of binding events throughout a single measurement. The decay constant can be determined by plotting number of detected particles for each species vs. time and fitting an exponential decay. We found that the resulting ratio of the decay constant for monomer:dimer is 1.04 $\pm$ 0.21 (N = 13 experiments) in line with expectations based on diffusion coefficient alone ($2^{1/3} = 1.26$), suggesting that our experiments are sampling monomer and dimer populations equally.
To obtain the binding plot from Fig. 1b in the main manuscript, we compared the tubulin system to the standard formula for a ligand receptor interaction[3]:

$$[RL] = \frac{[L][R_T]}{[L] + K_d} \tag{4}$$

where $[RL]$ is the concentration of ligand receptor complex, $[L]$ is the free ligand concentration, $[R_T]$ is the total receptor concentration. In our case, $[RL]$ is the concentration of monomers in the dimeric state, $[R_T]$ is the total monomer concentration and the $[L]$ is twice the free monomer concentration (due to R $\equiv$ L for a dimeric system), this equation becomes[4]:

$$2[D] = \frac{2[M][M]_{TOT}}{2[M] + K_d} \tag{5}$$

Dividing by the total monomer concentration and substituting for the corrected numbers of each species, the fraction of tubulin in the dimeric state becomes:

$$\frac{2cN_D}{[M]_{TOT}} = \frac{2cN_M}{2cN_M + K_d} \tag{6}$$

A plot of $\frac{2cN_D}{[M]_{TOT}}$ as a function $2cN_M$ can be fit to the following general logistic function:

$$f(x) = A_1 + (A_2 - A_1)\frac{x^n}{x^n + K_d^n} \tag{7}$$

$A_1$ and $A_2$ are the lower and higher asymptotes respectively and $n$ is the cooperativity of the association. In the high tubulin concentration limit, the fraction of tubulin in the dimeric state will be 1 so by setting $A_1$ to 0, $A_2$ to 1, and the cooperativity to 1, a single parameter least-squares fit was carried out to determine the value of $K_d$.

To calculate the $K_d$ values for each individual experiment (Fig. 1e), a direct substitution into equation 1 yields:

$$K_d = \frac{cN_M^2}{N_D} \tag{8}$$
